# Marchantia stem cell maintenance and re-establishment are controlled by MpPIN1-mediated auxin transport and ARF signalling

**DOI:** 10.64898/2026.06.29.735252

**Authors:** Ignacy Bonter, Marius Rebmann, Mihails Delmans, Facundo Romani, Jim Haseloff

## Abstract

Meristem organization in land plants depends on coordinated auxin distribution, transport and response, yet how these processes are integrated during vegetative growth of the gametophyte of non-seed plants remains poorly understood. Here, we describe dissection of the auxin network in the liverwort *Marchantia polymorpha* using a suite of endogenous and synthetic reporters. Quantitative imaging with the R2DII auxin response sensor revealed that the apical meristem is positioned at a stable auxin minimum, with auxin levels increasing toward differentiated tissues. This spatial configuration correlates with polar localization of the auxin efflux carrier MpPIN1 along the direction of auxin flux. Disruption of Mp*PIN1* alters auxin distribution and compromises tissue regeneration, but does not prevent initial meristem formation, indicating that PIN-mediated transport reinforces rather than defines stem-cell niche identity. Imaging of auxin response reporters and Mp*ARF1* and Mp*ARF2* knock-in reporters further revealed a constrained auxin response within the meristem, characterized by low auxin signalling, dependent on the inhibitory activity of Mp*ARF2* in the stem cell zone. During regeneration, transient, oscillatory *ARF* dynamics were observed before stabilization of a new auxin minimum and re-establishment of normal meristem architecture. Together, our results are consistent with auxin responses in the Marchantia meristem being shaped by an incoherent feed-forward network topology that couples auxin flux, transcriptional constraint and niche permissiveness. This architecture provides a robust framework for stem-cell maintenance and re-establishment, revealing ancestral principles of meristem regulation in land plants.

**SUMMARY STATEMENT:** By mapping hormone distribution at cellular resolution in the liverwort *Marchantia polymorpha*, we reveal fundamental principles governing how plants maintain and rebuild their stem cell niches.

## INTRODUCTION

The liverwort *Marchantia polymorpha* has emerged as a powerful model in plant evolutionary developmental biology (evo-devo). Its position among the earliest-diverging lineages of land plants makes it an ideal reference for reconstructing ancestral mechanisms of plant development. At the same time, Marchantia offers practical advantages: it has a compact, non-duplicated genome, minimal genetic redundancy, and single-copy representatives of key developmental regulators and hormonal pathway components (Bowman et al., 2017; Blazquez et al., 2020). In contrast to the sporophyte-based meristems of flowering plants such as *Arabidopsis thaliana*, the Marchantia meristem operates in the gametophyte generation, providing a unique opportunity to explore the evolution of stem cell regulation.

The vegetative meristem of Marchantia is located in a dorsal indentation known as the apical notch. Recent studies have provided exceptional details about the cellular architecture of the gametophyte meristem and have refined this view to a discrete stem cell zone centred on a small population of apical cells (Romani et al., 2024; Spencer et al., 2024). This apical cell lies subapically on the ventral side of the thallus and divides from four cutting faces to generate the tissues of the dorsal and ventral thallus. The apical cell and its immediate progeny form the stem cell niche, surrounded by progressively differentiating cells.

Visualizing the meristem at cellular resolution remains technically challenging, as the apical region is covered by pigmented scales and chlorophyll-rich air chambers. Consequently, most image analysis derives from histological sections or studies in early development, when the mature meristem is not yet established. Lineage-tracing and clonal analyses have established that positional cues, rather than cell lineage, determine developmental patterns (Suzuki et al., 2020), reinforcing the view of the Marchantia meristem as a spatially organized niche centred on a single, self-renewing apical cell.

The single apical cell meristem is considered ancestral for land plants. In contrast, flowering plant meristems have evolved multicellular stem cell niches maintained by complex feedback circuits, most notably the *WUSCHEL–CLAVATA* (*WUS–CLV*) pathway. In our previous work (Romani et al., 2024), we established a transcriptional map of the Marchantia vegetative meristem. By integrating spatial transcriptomics and reporter analyses, we identified transcription factors (TFs) enriched in specific meristematic domains. Comparative analysis revealed striking differences between the Marchantia gametophyte and the Arabidopsis sporophyte, including the functional divergence of classical meristem regulators such as *KNOX* and *WOX* family members (Hirakawa et al., 2020; Hisanaga et al., 2021; Romani and Moreno, 2021). These findings indicate that gametophyte meristems are governed by distinct genetic circuits.

Among the TFs we identified, Mp*ERF20* (also known as *LOW-AUXIN RESPONSIVE*, Mp*LAXR*), an ethylene response factor, is specifically expressed in the apical cell and is rapidly induced during regeneration (Romani et al., 2024). Functional analyses revealed that Mp*ERF20* contributes to new meristem formation following wounding, linking it to the establishment of stem cell identity (Ishida et al., 2022). Marchantia’s remarkable regenerative capacity offers an opportunity to study de novo meristem formation. Upon wounding, a transient drop in auxin precedes the induction of Mp*ERF20* (Ishida et al., 2022). Exogenous auxin application inhibits regeneration and suppresses Mp*ERF20* expression, suggesting that low auxin levels are required to reinitiate stem cell programs (Nishihama et al., 2015; Ishida et al., 2022). This interplay between auxin depletion and transcriptional reprogramming highlights the tight hormonal control of the processes required for regeneration.

Auxin is a pivotal regulator of plant morphogenesis. The nuclear auxin signaling pathway depends on TIR1/AFB receptor for hormone perception. In Marchantia, Mp*tir1* mutants form disorganized cell masses lacking organs or meristems (Suzuki et al., 2022), demonstrating that auxin signalling is essential for normal organogenesis. Auxin biosynthesis proceeds through the indole-3-pyruvic acid (IPA) pathway, involving the enzymes MpTAA and MpYUC2, both expressed in the apical notch and gemma cups (Eklund et al., 2015). Apical localization suggested that auxin is synthesized close to the meristem; however, whether auxin forms a gradient, accumulates in specific cell layers, or how it is dynamically modulated during regeneration has not been experimentally resolved.

Directional transport of auxin is largely mediated by *PIN-FORMED* (*PIN*) efflux carriers. Marchantia possesses a single full-length *PIN*, Mp*PIN1*, which localizes to the plasma membrane and displays clear polarity in specific developmental contexts (Fisher et al., 2023). Mp*PIN1-GFP* polarity aligns with gravity and light cues in germinating gemmae and gametangiophores (Fisher et al., 2023), consistent with its role in polar auxin transport. Mp*pin1* mutants exhibit impaired phototropic and gravitropic responses and reduced regeneration efficiency but still form dichotomously branching meristems. These observations indicate that polar auxin transport refines auxin distribution and regeneration capacity but is not absolutely required for meristem initiation. It was recently shown that Mp*PIN1* is involved in the distribution of auxin in the gemma. (Flores-Sandoval et al., 2025). However, it is not clear how this auxin patterning is established and maintained in the mature meristem.

Auxin signalling output in Marchantia is governed by a minimal network of three *AUXIN RESPONSE FACTORS* (*ARFs*): the activator Mp*ARF1*, the repressor Mp*ARF2*, and the class C factor Mp*ARF3*. The latter, do not directly participate in auxin signalling (Flores-Sandoval et al., 2015; Kato et al., 2015; Kato et al., 2017; Flores-Sandoval et al., 2018; Kato et al., 2020). The ratio between MpARF1 and MpARF2 determines the cellular sensitivity to auxin (Kato et al., 2020) and it is tightly controlled by protein degradation (de Roij et al., 2025). High MpARF2 and low MpARF1 levels in apical cells create an auxin-insensitive zone, preventing premature differentiation even in the presence of auxin synthesis nearby. It has been shown that Mp*ARF2* plays a critical role in meristem maintenance and establishment (Flores-Sandoval et al., 2025). The spatiotemporal coordination between auxin distribution, ARF activity, and transcriptional regulators such as Mp*ERF20* therefore represents a key frontier in understanding meristem function.

Despite extensive molecular and genetic evidence that auxin signalling is critical for development in Marchantia, the precise contributions of auxin dynamics and differential cellular responses to maintenance of the Marchantia meristem remain unclear. Here, we have systematically visualized a series of reporters, including an auxin R2DII reporter, to explore auxin distribution with cellular resolution in gametophyte development and regeneration. We reveal a stable auxin minimum in the apical region and a progressive increase in auxin response toward differentiating tissues. We also show that this gradient depends on MpPIN1, whose polar localization aligns with the direction of auxin flux. By integrating auxin reporter imaging, MpPIN1 polarity analysis, and auxin-responsive marker expression, we establish a mechanistic framework linking gene regulatory networks, hormonal signalling to stem cell identity. This work provides insight into the ancestral principles of auxin-mediated niche regulation in land plants.

## RESULTS

### Vegetative development is characterized by expansion of cellular domains during meristem maturation

During gemma germination, the division and differentiating cell zone (DDCZ) within the apical notch expands from a restricted domain in the gemma to a broad gradient in the mature thallus (Romani et al., 2024). The adult Marchantia mature meristem is defined by active cell division (Romani et al., 2026) and the production of air pores dorsally and scales ventrally, which begin to develop after a series of cell divisions 3-4 days after gemma germination.

Previously, we generated a series of reporter lines driven by promoters of TFs expressed in distinct regions of the gemma apical notch (Romani et al., 2024). To further characterise TF expression in the mature meristem, we selected a combination of three markers, with p*ro*Mp*BZIP9*, pr*o*Mp*BZIP7*, and *pro*Mp*ERF7* each driving expression of a fluorescent protein. The expression domains showed a nested arrangement around the apical notch. To follow developmental transitions, we imaged both gemmae and 5-day-old gemmalings expressing the markers, during the time when the meristem becomes fully established.

All three reporters expanded their expression domains during early development while maintaining their relative spatial order: *pro*Mp*BZIP7* drives expression closest to the notch, following by *pro*Mp*ERF7* and *pro*Mp*BZIP9* marking the more distal cells. In the 5-day old thallus, *pro*Mp*BZIP7* is restricted a region within the DDCZ, while proMpBZIP9 covers all the DDCZ, and *pro*Mp*ERF7* is active in the DDCZ and beyond (Figure 1A,C).

**FIGURE 1.**
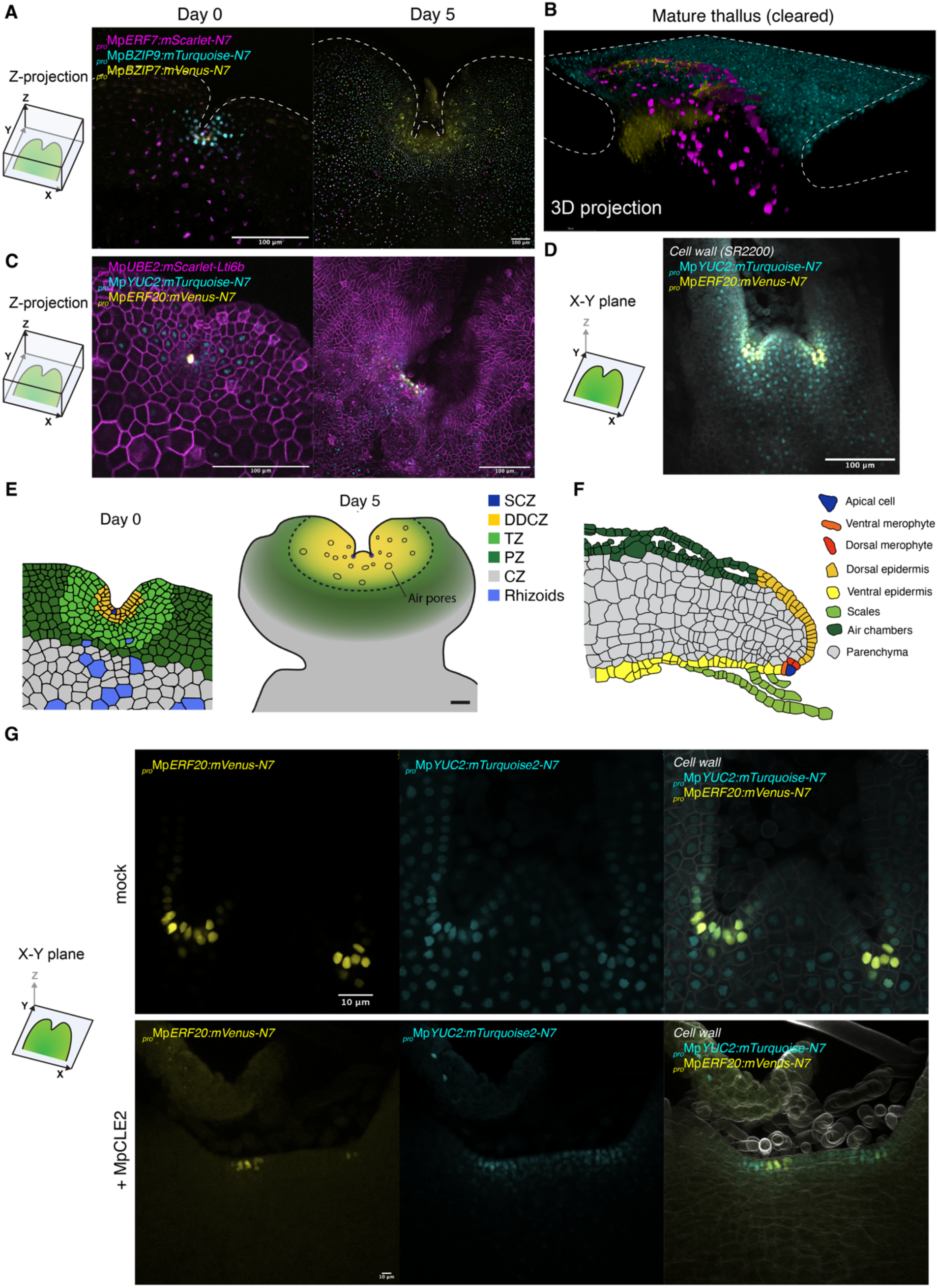
Expansion of expression domains in gemmalings. (A) Z-stack confocal images of transcription factor promoters of plants transformed with proMpERF7:mScarlet-N7 (magenta), proMpBZIP7:mVenus-N7 (yellow), proMpBZIP9:mTurquoise-N7 (cyan) reporters in a gemma and 5-day-old gemmaling. (B) 3D projection of two-week-old plants optically cleared of the same reporter as A. (C) Z-stack confocal images of plants transformed with proMpERF20:mVenus-N7 (yellow) and proMpYUC2:mVenus-N7 (cyan), and proMpUBE2:mScarlet-Lti6b (magenta) reporters in a gemma and 5-day-old gemmaling. (D) Confocal image of the X-Y plane centred in the apical notch of optically cleared plants with the same reporter as in C in 2-week-old plants. (E) Diagram of expression domains and cell types found in gemmae and 5 days gemmalings. (F) Diagram of cell types from a dorsal view of the thallus specifying cell types. (G) Confocal image of the X-Y plane centred in the apical notch of optically cleared plants expressing the same reporter as in C. Plants grown for 7 days in either normal media (mock) or supplemented with 30 μM MpCLE2 peptide. Scale bars are shown labelled in each panel.

As the gemmalings mature, it is difficult to visualize the apical cells, as they become obscured below the thallus (Figure 1B,D). To visualise expression in the mature thallus, we applied optical clearing to image deeper cell layers (Sakamoto et al., 2022). In two-week-old plants, *pro*Mp*BZIP7* expression remained associated with dividing cells around notch; *pro*Mp*BZIP9* was confined to the upper epidermis, air chambers, and photosynthetic filaments; and *pro*Mp*ERF7* became specific to the scales covering the meristem. This pronounced reorganisation underscores the significant transitions that the meristem undergoes as the thallus matures.

Among meristematic markers, *pro*Mp*ERF20/LAXR* and *pro*Mp*YUC2* have been used to label meristematic domains, although their spatial patterns have not been directly compared. We generated a dual-reporter line combining both reporters. Confocal imaging of cleared meristems confirmed that *pro*Mp*ERF20* marks the SCZ, apical and immediate sub-apical cells in the mature meristem, showing a narrower expression domain than *pro*Mp*YUC2*, which defines the DDCZ.

Ectopic or exogenous application of the peptide MpCLE2 has been shown to perturb meristem organization, resulting in an increase in the number of apical cells in Marchantia (Hirakawa et al., 2020). This expansion was previously seen in anatomical observations (a wider notch with more cells) and in an enlarged *pro*Mp*YUC2:GUS* domain following Mp*CLE2* overexpression or peptide application. To explore how these treatments affect cell identity, we analysed the expression of both proMpERF20 and *proMpYUC2* after MpCLE2 peptide treatment. Our results confirm the expansion of the apical notch and Mp*YUC2* expression domain. However, despite the altered morphology, two discrete centres of Mp*ERF20* expression remained visible (Fig 1G). This indicates that MpCLE2 application alters the anatomy of the apical notch, potentially by inhibiting cell divisions in the middle lobe. This results with the arrest of subsequent branching events, consistent with (Hirakawa et al., 2020).

### An auxin gradient is formed around the low auxin zone at the apical notch

It has been proposed that the apical notch in the gametophyte of Marchantia contains a zone with low auxin levels (Flores-Sandoval et al., 2025; Wallner et al., 2026) that this regulates Mp*ERF20* expression in the SCZ (Ishida et al., 2022). However, the spatial distribution of auxin is not very clear, particularly in the mature apical meristem. To investigate this, we generated an R2DII ratiometric reporter adapted for Marchantia *(Liao et al., 2015)*.

The R2DII system relies on the auxin-dependent degradation of the DII motif of Aux/IAA proteins through the TIR1-mediated ubiquitination pathway, and it is widely used to visualize *in vivo* auxin concentrations. In our design, the reporter comprises two nuclear-localized fluorescent proteins: one containing an intact DII motif (DII–mVenus–N7) and another carrying a mutated, non-degradable DII variant (mDII–mTurquoise2–N7). Both are expressed as independent transcriptional units under the control of the constitutive Mp*UBE2* promoter. The point mutation in the DII domain prevents interaction with TIR1 in the presence of auxin, thus stabilizing the fluorescent signal. Consequently, the mVenus/mTurquoise fluorescence ratio reflects intracellular auxin concentration independently of downstream auxin response factors.

To validate R2DII in Marchantia, plants were treated overnight with 50 µM NAA. The reporter responded as expected, showing decreased mVenus–N7–DII fluorescence relative to mTurquoise2–N7, confirming its sensitivity to auxin. Using this system, we detected an auxin gradient with a minimum in the stem cell zone (SCZ) and a maximum in the central zone (CZ) and mature thallus (Figure 2A,B). This pattern is more prominent in the mature thallus and consistent with the proposed auxin minimum in the apical cells inferred from the auxin-dependent expression of proMpERF20/LAXR (Ishida et al., 2022). A similar auxin minimum was also evident in the midrib of the ventral epidermis (Figure 2C).

**FIGURE 2.**
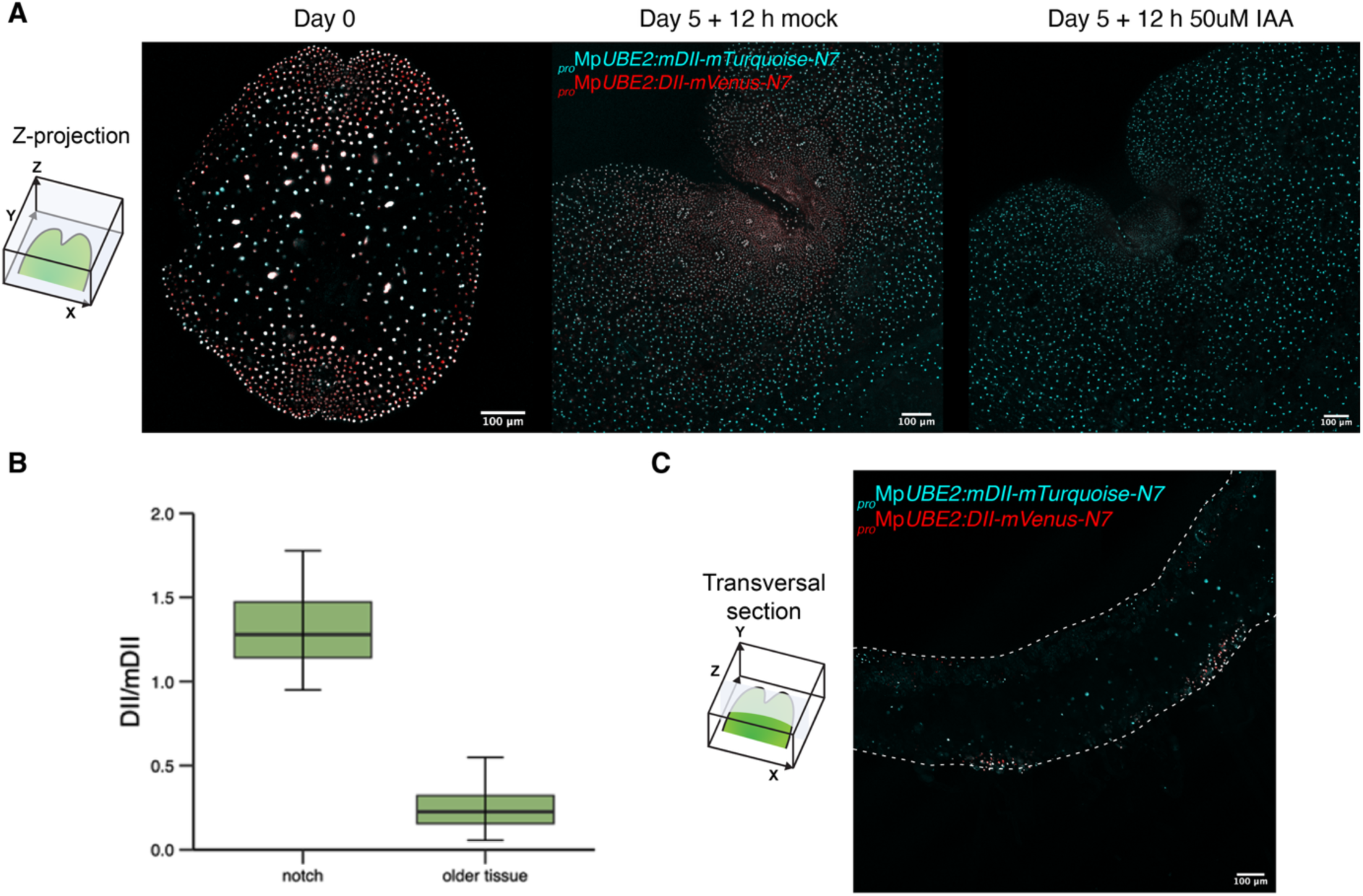
Spatial distribution of auxin in gemmalings. (A) Z-projection confocal image of plants transformed with the R2DII ratiometric auxin reporter (proMpUBE2:mDII-mTurquoise-N7; proMpUBE2:DII-mVenus-N7) in gemma and in 5-day-old gemmalings treated with 50 µM IAA and imaged after 12 hours. Red regions indicate lower auxin accumulation. (B) DII-mVenus/mDII-mTurquoise pixel intensity ratio was calculated by average intensity in nuclear cross-section (high values indicate less auxin accumulation, n > 80). Asterisks indicate statistical significance, t-test (two-sample assuming unequal variances) p-value = 6.578 x 10^-82^. (C) Transverse section of the R2DII reporter in 2-week-old plants. Scale bars = 100 µm.

During sporeling development, the meristem is formed *de novo* from a single cell spore, as evidenced by the expression of *pro*Mp*ERF20* in the prothalloblast stage (Wallner and Dolan, 2024; Wallner et al., 2026). We generated spores for the R2DII reporter to establish when the auxin gradient is formed. After the first cell division, it is clear that the rhizoid has higher auxin concentration. In the young flabellum, there is a relatively low auxin concentration across the tissue, but the gradient is only visible after the notch with meristem is formed in ∼7 days old sporelings. After the maturation of the thallus at ∼10 days, the auxin minima region expands and converges with the pattern observed in the thallus of older gemmalings (Figure 3).

**FIGURE 3.**
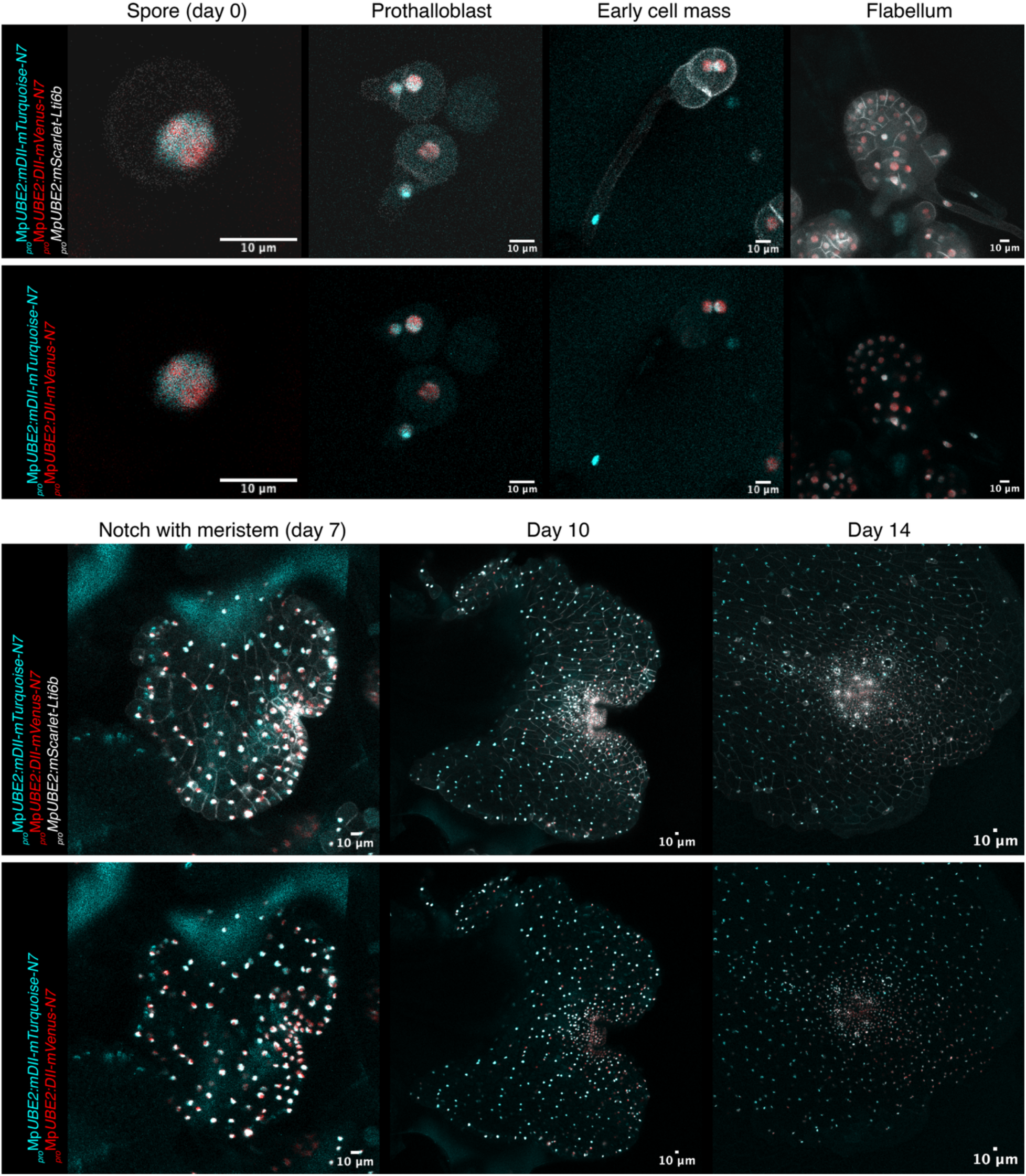
Spatial distribution of auxin in sporelings. Z-projection confocal images of plants transformed with the R2DII radiometric auxin reporter (proMpUBE2:mDII-mTurquoise-N7; proMpUBE2:DII-mVenus-N7) in a time series of sporelings imaged over 0-14 days. Representative pictures of developmental stages are shown. Red regions indicate lower auxin accumulation. Scale bars = 10 um.

**FIGURE 4.**
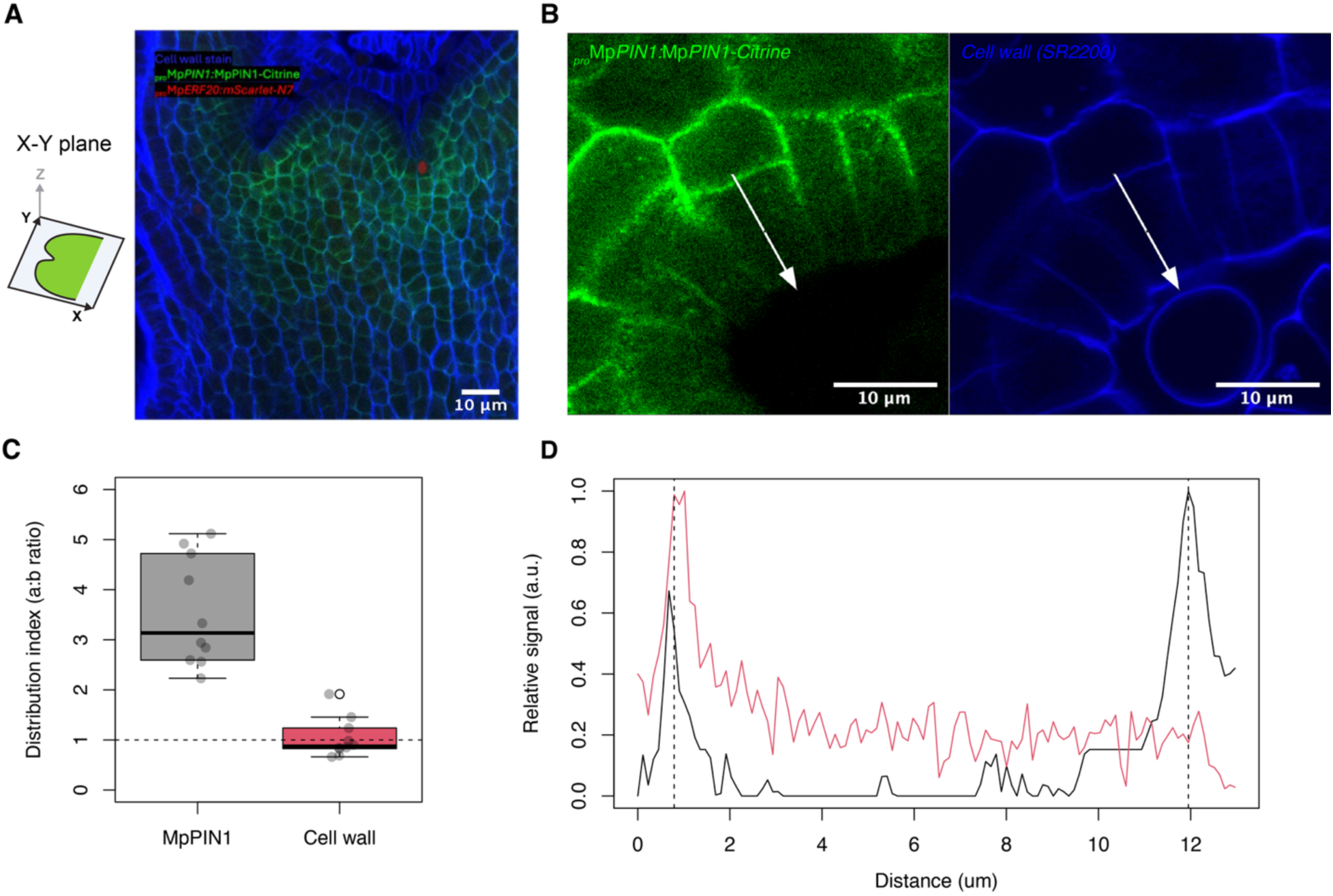
MpPIN1 protein is polarized towards the basal part of the thallus in the mature meristem. (A) Confocal image of the X-Y plane centred in the apical notch of a translational reporter of MpPIN1 (proMpPIN1:MpPIN1-Citrine, green) in the Mppin-1 background (Fisher et al., 2021) of 2-week old sporelings. The cell wall was stained with SR2200 (blue). (B) Close up of the apical notch. (C). Pixel intensity measurements of fluorescence of cell wall and MpPIN1 in opposite cells wall in the SCZ (n=10). (D) Quantification of fluorescence intensity (arbitrary units) a single cell corresponding to the arrow in B. Scale bars = 10 µm.

**FIGURE 5.**
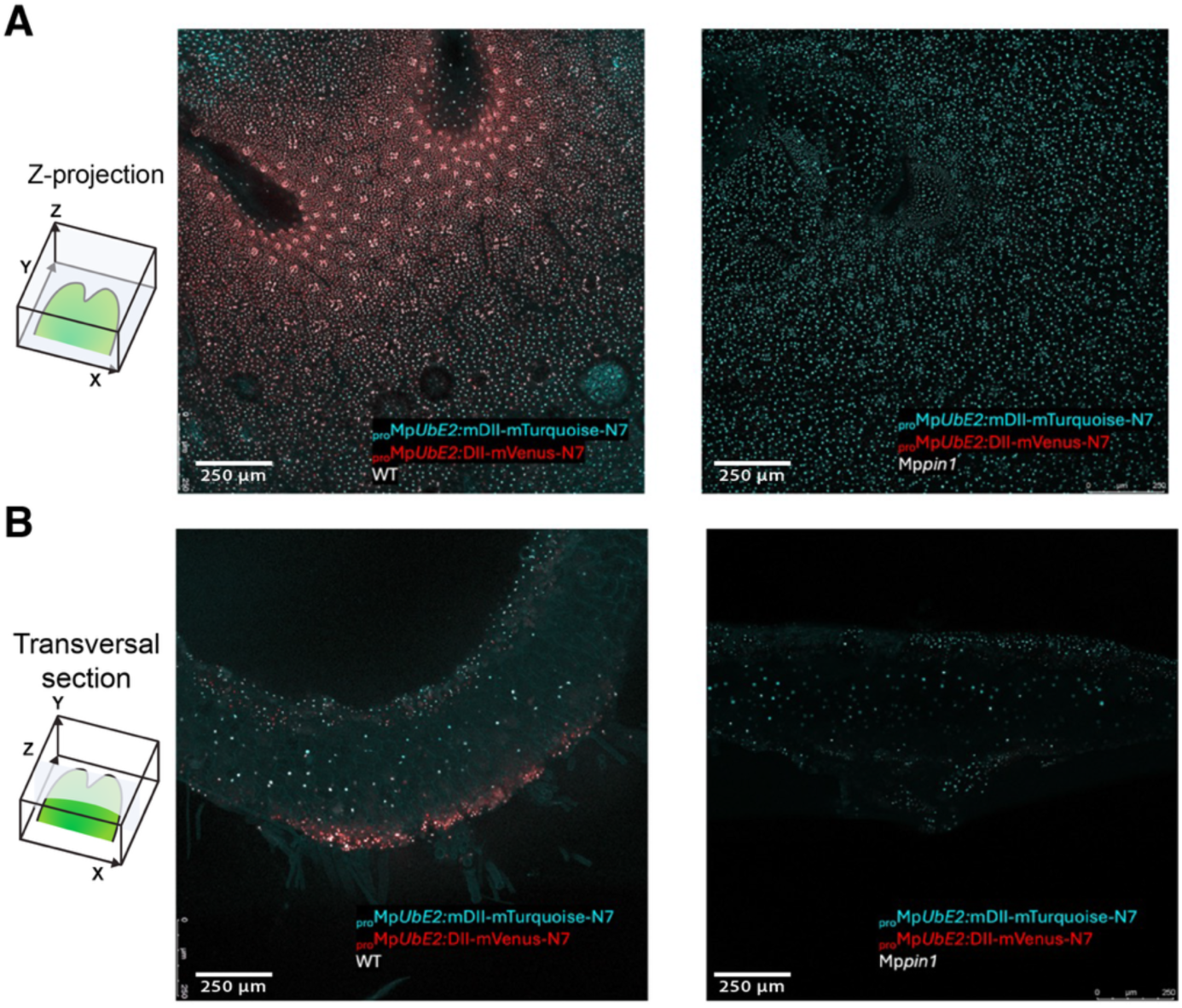
R2DII in the Mppin1 mutant background. (A) Z-axis projection of confocal images of the R2DII ratiometric auxin reporter in the WT and Mppin1 mutant in 4-weeks old Cam-1 plants. Red regions indicate lower auxin accumulation. (B) Transverse section of the same plants. Scale bars = 250 um.

### MpPIN1 mediated transport is required for the formation of the auxin gradient

The progressive establishment of the auxin gradient during sporeling maturation raised the question of which transport mechanisms are responsible for its formation and maintenance in the vegetative meristem. In flowering plants, polar auxin transport mediated by PIN1 is essential for establishing auxin gradients. In *Marchantia polymorpha*, MpPIN1 is the only canonical PIN protein with a predicted role in intercellular auxin transport (Fisher et al., 2023). To investigate its distribution and polarity in the adult meristem, we generated double transgenic lines by co-transforming two constructs conferring resistance to either hygromycin or G418, carrying the markers *pro*Mp*PIN1:*Mp*PIN1–Citrine*, (Fisher et al., 2023), and *pro*Mp*ERF20:mScarlet*.

In mature meristems, Mp*PIN1* expression expanded to a broader domain encompassing several layers of cells within the meristem. In cells around the apical notch, the MpPIN1–Citrine signal exhibited clear subcellular polarity. Co-staining with a cell wall marker confirmed that MpPIN1 accumulated on the basal side of the cells, consistent with basipetal export of auxin. While PIN polarity has been previously reported during gametangiophore development (Fisher et al., 2023) and gemma dorso-ventralization, this is the first demonstration of apico-basal PIN1 localization within the vegetative meristem.

To determine whether polar auxin transport contributes to the auxin gradient observed in the apical notch, we introduced the R2DII ratiometric reporter into an Mp*pin1* mutant background. In these plants, the auxin minimum in the apical notch was abolished, and a more uniform auxin distribution was observed (Fig.5). These results confirm that MpPIN1-mediated transport is required for establishing the auxin gradient in the Marchantia meristem.

### Auxin insensitivity in the stem cell zone reinforces the auxin minima

Auxin signalling output is determined not only by auxin concentration but also by the antagonistic activity of two *AUXIN RESPONSE FACTORS* (*ARF*) TFs (Mp*ARF1* and Mp*ARF2*) (Kato et al., 2020; Das et al., 2024; de Roij et al., 2025). Distinct cell types in Marchantia exhibit contrasting auxin sensitivities: the stem cell zone (SCZ) is characterized by high Mp*ARF2* expression and low auxin responsiveness, whereas rhizoid cells show elevated Mp*ARF1* levels and strong auxin sensitivity (Das et al., 2024). The spatial pattern of auxin sensitivity therefore parallels the auxin concentration gradient, suggesting that both phenomena act in concert to reinforce each other. This expression pattern is particularly true at the gemma stage and such gradient is maintained in the mature apical notch (Figure 6A). Interestingly, elevated MpARF2/MpARF1 ratio is also prominent in developing air pores (Figure 6B).

**FIGURE 6.**
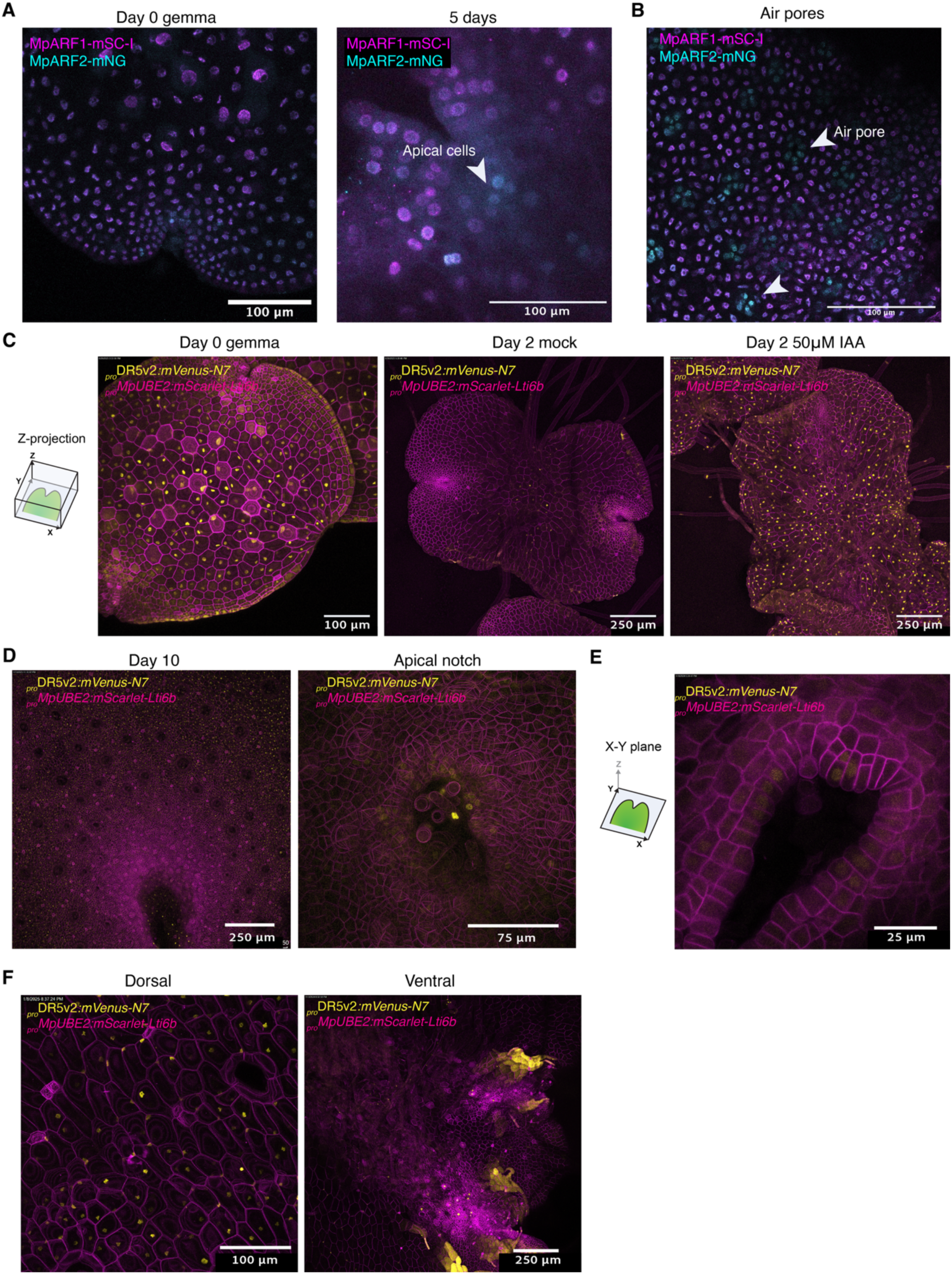
Auxin Responsive Factor expression and auxin signalling in the thallus. (A-B) Optical projections of Z-axis confocal images of knock-in lines for MpARF1 and MpARF2 fused to mScarlet-I (magenta) and mNeon Green (cyan) fluorescent proteins, respectively (Das et al., 2024). Gemmae and 3-day-old and 1-week gemmalings are shown. Arrows highlight developing air pores. (C) Projected Z-stack confocal images of proDR5v2:mVenus-N7 (yellow) in gemmae and plants grown for 2-days in either mock or 50 µM NAA. The latter two have the same laser and image settings. (D) Projected Z-stack confocal images of proDR5v2:mVenus-N7 in 10 day-old gemmalings after the establishment of the mature thallus (left). A close-up look of the apical notch is shown (right). (E) Confocal image of the X-Y plane centred in the apical notch of proDR5v2:mVenus-N7 in 2-week-old plants. (F) Z-stack confocal images of the dorsal (left) and ventral (right) sides of 2-week-old proDR5v2:mVenus-N7 plants. In C-F, a constitutive plasma membrane marker is also shown (magenta, proMpUBE2:mScarlet-Lti6b). Scale bars are labelled in each panel.

To visualize auxin response activity, we employed the synthetic *proDR5v2* reporter, which contains ARF-binding sites and serves as a proxy for the transcriptional output of auxin signalling (Liao et al., 2015), downstream of ARFs. The DR5 signal was strongest in older tissues distant from the notch, where cells had completed expansion and differentiation, likely functioning as an auxin sink (Figure 6C-D). Notably, while the dorsal epidermis displayed prominent DR5 activity, neither air pore cells nor oil body cells showed detectable signal. This is consistent with the pattern of auxin distribution shown by the R2DII reporter and the protein levels of MpARF1/2 (Das et al., 2024). In contrast, the ventral epidermis generally lacked signal, except for rhizoid precursor cells, which showed the strongest DR5 fluorescence of all tissues examined (Figure 6E).

To resolve patterns of auxin response within the meristem, we performed chemical fixation and optical clearing of mature thalli. This revealed distinct DR5 expression in cells flanking the apical cell and at the primordium of the emerging middle lobe (Figure 6D), suggesting that auxin signalling is not limited to mature cells, but also local activity could be important for organ formation adjacent to the central part of the meristem.

Overall, these results are consistent with the pattern of auxin distribution showed by the R2DII reporter and the protein levels of MpARF1/2 (Das et al., 2024). This supports a model in which differential auxin sensitivity, driven by opposing MpARF1 and MpARF2 activity, establishes a domain of auxin insensitivity in the SCZ, reinforcing the auxin minimum at the apical notch. The same processes seem to govern gemmae formation within the cup structure, as we found evidence of active auxin transport, high MpARF2 levels, low auxin concentration and response levels within the gemmae primordia (Supplementary Figure 1).

### Auxin transport and signalling contribute to the re-establishment of the stem cell niche during regeneration

Our data so far describe how auxin biosynthesis, transport, and signalling interact to maintain meristem homeostasis. However, the auxin gradient appears to be strictly required for vegetative growth (Fisher et al., 2023), as revealed by the R2DII reporter in the Mp*pin1* mutant. To test how these components act in the context of meristem re-establishment, we analysed apical cell regeneration following notch excision.

Auxin plays a central role in regeneration in *Marchantia (Nishihama et al., 2015)*. Removal of the apical notch triggers a sharp drop in auxin concentration (Ishida et al., 2022), likely due to the loss of local biosynthesis. This is accompanied by an induction of Mp*ERF20/LAXR* after 6 hours (Ishida et al., 2022; Romani et al., 2024). Consistent with this, Mp*YUC2*, the only *YUCCA* gene expressed in the gametophyte, and MpPIN1 are also upregulated after 6 and 12 hours respectively (Figure 7A). On the other hand, Mp*ARF2* expression increases and Mp*ARF1* decreases (Figure 7A). The series of early events taking place during regeneration, possibly start with a sharp increase of Mp*ARF2* ratio, to increase auxin insensitivity; followed by Mp*ERF20/LAXR* and Mp*YUC2* activation, to restore meristem identity and auxin biosynthesis; and finally, possible export of auxin from the notch via Mp*PIN1*.

**FIGURE 7.**
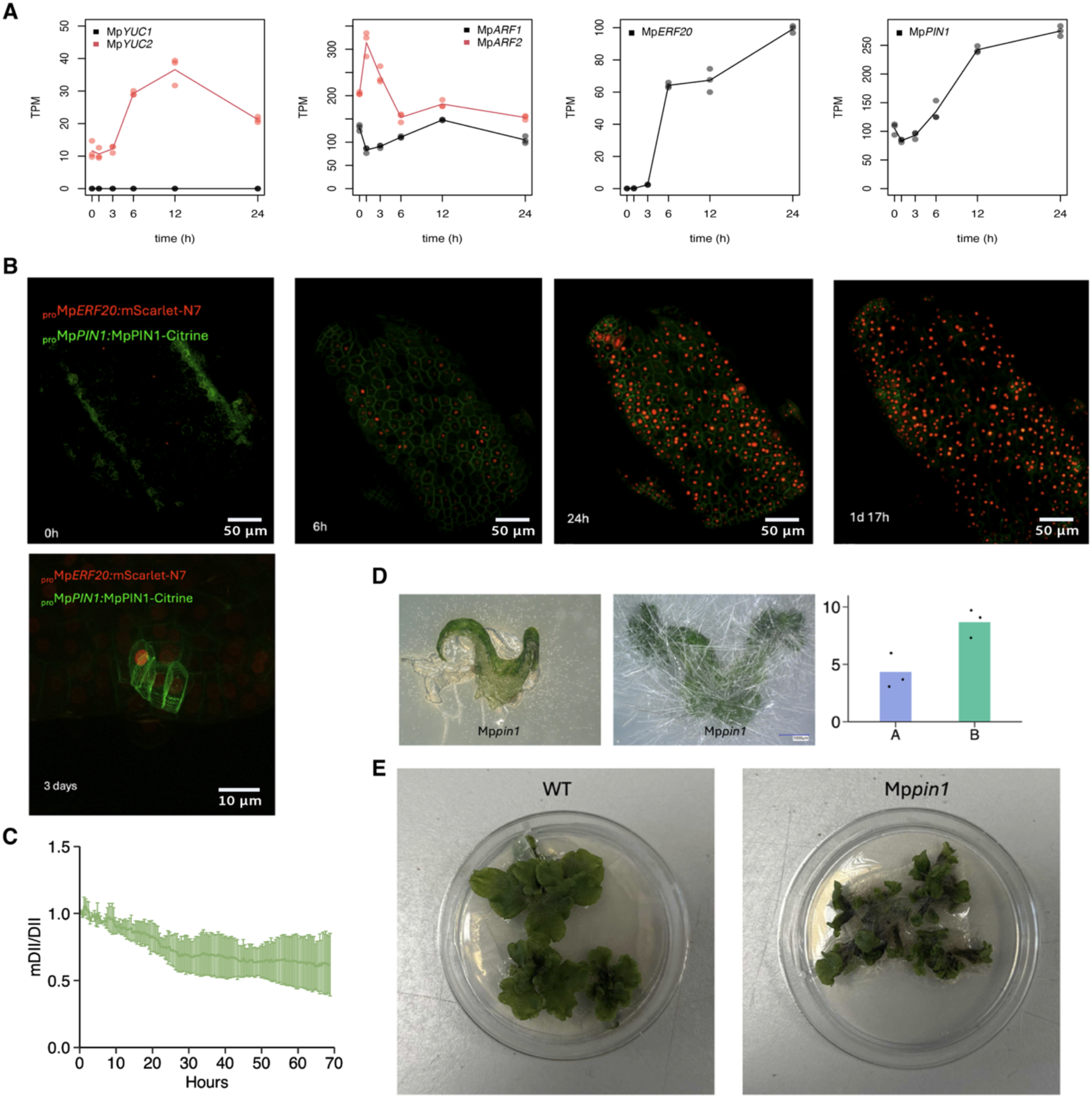
Gene expression dynamics during meristem re-establishment. (A) RNA-seq analysis of representative cell cycle genes during the time course of basal portion of regenerating plants (Ishida et al., 2022). Genes are grouped by families and distinguished by colour (see legend). Individual points represent biological replicates and lines show average values. (B) Top row, time course images of the regenerating central portion of a gemma expressing proMpERF20:mScarlet-N7 and proMpPIN1:MpPIN1-mCitrine, bottom panel, close up image of new apical cells three days post wounding. (C) Quantification of average mDII/DII fluorescence ratio in three regenerating buds over 70 hours of continuous imaging during regeneration. (D) Excised thallus fragments of Mppin1 plants 0 (left) and 5 (centre) days after wounding. Right, quantification of new meristems in WT and Mppin1. (E) Overview of regenerating plant fragments two weeks after wounding in WT and Mppin1.

To visualise auxin transport during this process, we imaged regenerating thalli of the Mp*pin1*^ge^ genetic complementation line co-expressing *pro*Mp*PIN1:*Mp*PIN1–mCitrine* and *pro*Mp*ERF20:mScarlet–N7*. Time-lapse imaging over three days revealed two distinct phases. Initially, Mp*PIN1* and Mp*ERF20* were induced broadly and appeared non-polar in the membranes of all central cells. By 24 hours, Mp*PIN1* translational reporter became restricted to regenerating buds with high cell division activity, where it gradually acquired a polar distribution. By day three, Mp*PIN1* persisted only in apical and subapical cells, matching the domain of Mp*ERF20* promoter activity (Romani et al., 2024), whereas surrounding tissue showed diluted signal (Figure 7B, Suppl. Movie 1).

Previous work described a delayed regeneration phenotype in Mp*pin1* mutants (Fisher et al., 2023). We confirmed this by quantifying meristem initiation in regenerating explants. Although Mp*PIN1* was not required for the initial reprogramming of differentiated cells, mutants formed more numerous but smaller outgrowths, consistent with prolonged maintenance of undifferentiated callus-like tissue (Figure7D-E). This suggests that MpPIN1-mediated auxin transport contributes to apical dominance during regeneration, limiting the number of new growth foci and preventing self-shading or resource competition.

To test whether this effect correlates with auxin distribution, we performed time-lapse imaging of R2DII reporter plants regenerating for three days (Figure 7C). Quantification of the mDII/DII ratio within newly formed buds revealed a ∼40% reduction over time, consistent with the re-establishment of an auxin gradient dependent on MpPIN1 activity. However, it should be noted that R2DII ratio may not be able to capture rapid changes in auxin concentration during regeneration.

We next examined the dynamics of auxin signalling components. Time-lapse imaging of MpARF1–mScarlet and MpARF2–mNeonGreen knock-in lines (Das et al., 2024) revealed a transient accumulation of both proteins in nuclei during the first 6–8 hours after wounding, followed by rapid degradation. This early burst of ARF accumulation corresponds with transcriptomic evidence of wound-induced expression, although the absence of detectable MpARF2 protein immediately after wounding suggests translational or post-translational control. We propose that the initial spike in ARF abundance reflects a temporary inhibition of *ARF* degradation to rapidly modulate auxin sensitivity.

To assess the functional impact of these changes, we imaged regenerating DR5v2 reporter plants over 60 hours (Figure 8A, Suppl. Movie 2). Non-regenerating tissues exhibited a progressive increase in DR5-driven mVenus signal, indicating elevated auxin response, whereas regenerating buds lacked detectable signal. This pattern is consistent with a high MpARF2/MpARF1 ratio in the regenerating cells (Figure 8B), where inhibitory ARF2 proteins likely compete with the activator ARF1 for binding to auxin-responsive elements, maintaining local auxin insensitivity required for apical identity.

Interestingly, quantitative analysis of MpARF2–mNeonGreen fluorescence revealed rhythmic oscillations in abundance of the nuclear-targeted reporter proteins during regeneration, with a period of approximately 5–6 hours (Figure 8C-D). These oscillations were synchronized among buds within a single thallus but not across independent regenerating plants, suggesting the presence of a systemic, but plant-specific, oscillatory signal distinct from circadian regulation.

Together, these results indicate that MpPIN1-mediated transport and dynamic *ARF* regulation together facilitate the reconstruction of a low-auxin, auxin-insensitive niche, allowing the re-emergence of apical cell identity during regeneration.

## DISCUSSION

In this work, we provide a comprehensive and detailed view of the auxin signalling networks governing meristem development in the gametophyte of Marchantia. The set of known and novel reporters presented here, provide cellular-scale detail for a consistent model of stem cell maintenance (Figure 1-2), *de novo* formation of meristems in sporelings (Figure 3) and regeneration after wounding (Figure 7-8). Our results demonstrate that Marchantia employs a closely linked auxin signalling architecture, with neighbouring cells displaying distinct patterns of auxin synthesis, transport and response that establish critical spatial patterns within the vegetative gametophytic meristem. This type of inter-relationship parallels the situation in angiosperm sporophytes, where dynamic structures such as the shoot apical meristem contain auxin-insensitive apical cells, while *DR5* signals are found concentrated in emerging organ primordia (Ulmasov et al., 1997; Heisler et al., 2005; Vernoux et al., 2010).

**FIGURE 8.**
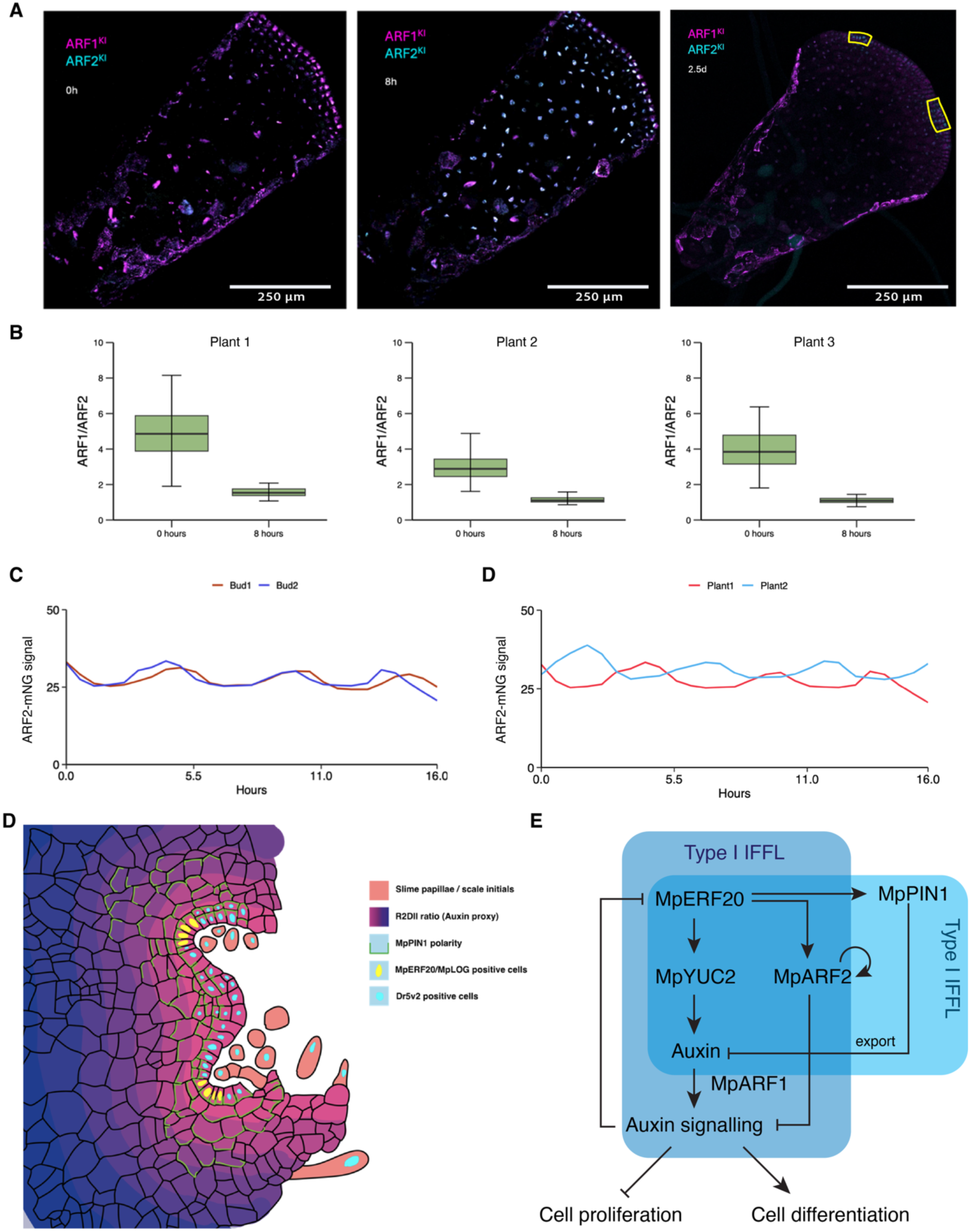
(A) Projected Z-stack images of the central portions of regenerating gemmae at 0, 8 and 36 hours with MpARF1 and MpARF2 fused to mScarlet-I (magenta) and mNeon Green (cyan) fluorescent proteins, respectively. Yellow outlined regions on the last panel indicate regenerating buds with oscillating MpARF2 fluorescence. (B) Ratio of ARF1/ARF2 fluorescence at 0 and 8 hours after wounding (average values for all nuclei in the tissue in three gemmae). (C) Quantification of MpARF2-mNeonGree fluorescence in two regenerating buds in the same gemma fragment. Each plot correspond to independent biological replicates. (D) Proposed model of auxin signalling with blue indicating high auxin and magenta low auxin, superimposed on a model for key cell types in the central part of the meristem, with MpPIN1 localisation marked in green. (E) Proposed schematic model of the Type I Incoherent Feed Forward Loop (IFFL) for auxin signalling in Marchantia.

Yet, TFs regulating this process in the gametophyte and the sporophyte are broadly different (Romani et al., 2024). While in angiosperms *WUSCHEL* acts as an auxin response rheostat in the shoot apical meristem (Ma et al., 2019), in Marchantia this fundamental role is fulfilled by Mp*ARF2* (Kato et al., 2020; Flores-Sandoval et al., 2025). The organizational logic of a meristems characterised by auxin gradient governed by transcription factors that maintains stem cell identity, represents a fundamental mechanism for meristem regulation that could be conserved across land plants and predates the evolution of complex multicellular apical meristems in the sporophyte.

The functional conservation of *ERF* TFs from the *ENHANCER OF SHOOT REGENERATION* (*ESR*) clade as stem cell markers represents a particularly important conservation pattern. Mp*ERF20/LAXR* specificity to apical cells in Marchantia parallels the stem cell-specific expression of the orthologues in both *Physcomitrium patens* gametophytes (Hata et al., 2025) and angiosperm sporophytes (Kirch et al., 2003). In all land plants, *ESR* homologues are sufficient to generate new stem cells and are specially required for regeneration (Banno et al., 2001; Ishida et al., 2022). This conservation across gametophytic and sporophytic meristems suggests that ERF-mediated transcriptional control may have been present in the last common ancestor of all land plants.

There is no doubt of the fundamental contribution of auxin signalling to establish polar growth and organogenesis in the gametophyte (Suzuki et al., 2022). However, there is an apparent contradiction between the overlapping sites of Mp*YUC2* expression (marking auxin biosynthesis) and Mp*ERF20/LAXR* expression (which expression is dependent on low auxin). This highlights the importance of understanding auxin homeostasis at cellular resolution and the gene regulatory networks underlying it. In this work, we demonstrated that the domain of Mp*ERF20/LAXR* expression is much narrower than that of Mp*YUC2* (Figure 1). The apparent paradox is resolved through the coordinated regulation of auxin export (MpPIN1), and sensitivity (MpARF2), creating microenvironments where auxin production does not necessarily correlate with an auxin maxima and high auxin signalling, a principle that may apply broadly to hormone patterning in plant development.

Our findings suggest that auxin homeostasis within the Marchantia stem cell niche is best understood as a self-organizing system regulated by a Type I incoherent feed-forward loop (IFFL) (Mangan and Alon, 2003; Alon, 2007; Kaplan et al., 2008). In such type of motifs an input signal regulates a target gene through two opposing pathways. In the case of Marchantia, the transcription factors that are specific to the stem cell niche (MpERF20 and MpARF2) are simultaneously promoting cell differentiation (through auxin biosynthesis) and repressing cell differentiation (via auxin polar transport and lower auxin sensitivity), creating a dynamic balance that governs cell fate decisions. This regulatory architecture provides evolutionary advantages through noise filtering and enhanced system robustness, critical features for maintaining stem cell identity across developmental perturbations.

Specifically, we propose that two interconnected IFFL are operating in this network (Figure 8D-E). First, high levels of Mp*ERF20/LAXR* in the SCZ activate Mp*YUC2*, which is indirectly promoting auxin signalling, and Mp*ARF2*, which is repressing auxin signalling. Subsequently, auxin signalling will repress Mp*ERF20/LAXR*, in a negative feedback loop. This effect is balanced by a second IFFL where MpPIN1 is inhibiting auxin accumulation by exporting it to neighbour cells. In that sense, MpERF20/LAXR is repressing itself but in different cells, promoting lateral inhibition. Mp*ARF2* can also activates auxin biosynthesis and carry on the IFFL even without MpERF20/LAXR and it is strictly required for meristem maintenance (Flores-Sandoval et al., 2025). In that sense, active transport of auxin mediated by MpPIN1 (Figure 7) and rapid activation of MpERF20/LAXR are important to establish apical dominance during regeneration (Figure 7), but not essential afterwards. While more details in the interconnection of these nodes are still required, this architecture is well supported by the available evidence.

The oscillatory dynamics of MpARF2 protein levels observed during regeneration could represent transient disruptions of this equilibrium (Figure 8). The 5–6 hour periodicity, their coordination within individual plants, and lack of synchronization across plants are consistent with the behaviour expected from negative feedback loops embedded in feed-forward circuits (Mangan and Alon, 2003; Zhang et al., 2016). Such oscillations can be interpreted as the system’s search for a new stable state following injury-induced perturbation, converging onto steady spatial patterns of auxin distribution and sensitivity. This dual-layer control of auxin level and responsiveness may give the capacity for rapid response when perturbed and may help explain the capacity for the fast regeneration of liverworts like Marchantia. Similar auxin-related oscillations have been reported in Arabidopsis root meristems, where dynamic auxin responses pattern lateral root formation (Moreno-Risueno et al., 2010). The observation of oscillations in ARF protein abundance within an IFFL framework may provide a new insight into underlying mechanisms of control.

The formation of the mature meristem represents a critical developmental transition that establishes the complex three-dimensional organization necessary for sustained growth (Romani et al., 2024; Spencer et al., 2024). Our work reveals that auxin patterning is integral to this maturation process, with the establishment of stable gradients of auxin in the DDCZ (Figure 2-3), while auxin signalling is concentrated in mature cells that exit the cell division and differentiation zone (Figure 5). In developing tissues, such as sporelings, and early gemmalings, these gradients are not fully established until the meristem develops (Figure 3-4).

Our observations suggest that stem cells in Marchantia, similar to those in *P. patens*, exist within a low-auxin (Thelander et al., 2019) and potentially high-cytokinin microenvironment (Cammarata et al., 2023; Hata et al., 2025; Komatsu et al., 2025), as evidenced by apical cell specific Mp*LOG* expression (Supplement Figure 2), which supports the maintenance of apical cell identity. This hormonal signature appears conserved despite the distinct morphological contexts. PIN1 polarity in bryophytes confirms the ancestral nature of directional auxin transport mechanisms (Bennett et al., 2014; Viaene et al., 2014). Our demonstration of polar MpPIN1 localization within the Marchantia vegetative meristem extends this conservation to liverworts (Figure 4), establishing that PIN-mediated polar transport represents a shared bryophyte innovation that likely evolved early in land plant evolution.

However, roles of shared auxin-regulatory components show interesting divergences between moss and liverwort lineages. In Marchantia, MpPIN1 is critical for establishing basipetal auxin flow and maintaining the auxin gradient that defines the stem cell zone yet is dispensable for fundamental meristem maintenance and dichotomous branching (Fisher et al., 2023). This contrasts with the more central role of PIN proteins in moss leaf initiation and organ patterning (Bennett et al., 2014), suggesting that while the basic transport machinery is conserved, its integration into developmental programs has diverged significantly between bryophyte lineages.

While our work demonstrates that Mp*PIN1* contributes to local auxin gradient formation, the broader auxin transport landscape remains incompletely characterized. Recent work has revealed that Marchantia possesses multiple short PIN proteins in addition to the canonical long PIN1, some exhibiting plasma membrane localization and auxin export activity (Tang et al., 2026). This expanded *PIN* family, combined with potential roles for *AUX/LAX* influx carriers, *ABCB* exporters, and auxin conjugation systems (Barbez et al., 2012; Peret et al., 2012; Ranocha et al., 2013; Mellor et al., 2016), suggest that auxin transport involves more complex networks than previously appreciated.

Our work contributes to the growing understanding that stem cell niches employ similar organizational principles across diverse plants, despite substantial differences in morphology and life cycle organization. By revealing how a minimal auxin regulatory network can generate spatial complexity in a basal land plant lineage, our study bridges developmental patterning in bryophytes and the evolution of more complex meristems in vascular plants. These findings provide a framework for reconstructing the ancestral logic of plant stem cell regulation and its diversification during land plant evolution.

## ACKNOWLEDGMENTS

We thank Dr. Tom Fisher for sharing plasmids containing MpPIN1 constructs and Shubhajoit Das, Jan Willem Borst and Prof. Dolf Weijers for sharing the Mp*ARF1/2* knock-in line. This work was funded as part of the Biotechnology and Biological Sciences Research Council and Engineering and Physical Sciences Research Council grant Grant BB/L014130/1 for the OpenPlant Synthetic Biology Research Centre and the Biotechnology and Biological Sciences Research Council grant BB/T007117/1 to J.H. Biotechnology and Biological Sciences Research Council grant BB/F011458/1 for confocal microscopy. F.R. is a Leverhulme Early Career Fellow (ECF-2023-534) funded by the Leverhulme Trust and the Isaac Newton Trust (23.08(f)), I.B. is funded by the Herschel Smith Fund studentship.

## MATERIALS AND METHODS

### Plasmid construction

To generate new Level 0 parts, CDS and promoter regions from genes were extracted from *M. polymorpha Tak-1* genome version 5.1 (Bowman et al., 2017) genome and manually domesticated to remove internal BsaI and SapI sites using synonymous mutations for the CDS. L0 parts were synthesized by GENEWIZ following the standard syntax for plant synthetic biology with CDS and PROM5 or PROM and 5UTR overhangs and cloned into the plasmid pUAP1 (Addgene #63674) (Patron et al., 2015) by homology recombination. Other parts were derived from previous works (Sauret-Gueto et al., 2020; Romani et al., 2024) as specified in Suppl. Table S2. DNA parts were assembled into transcription units using Loop assembly (Sauret-Gueto et al., 2021). Four sets of plasmids were utilised for L1 and L2 cloning (*pCk1-4* and *pCsA-D*), as well as *pBy01* (Romani et al., 2024) or *pBy10* (Tse et al., 2024). Translational reporter of Mp*PIN1* and CRISPR gRNA constructs were described before (Fisher et al., 2021).

### Analysis of RNA-sequencing data

TPM values for sporeling germination and different developmental stages were extracted from Marpolbase Expression database (Kawamura et al., 2022) or downloaded from SRA for regeneration (DRR330148-DRR330173), mapped in M. polymorpha Tak-1 genome v5.1 using HISAT2 (Kim et al., 2019), ht-seq and EdgeR. Data was subsequently analysed with R to generate plots using customs scripts.

### Plant material and growth conditions

*Marchantia polymorpha* subs*. rudelaris* accessions *Cam-1* (male) and *Cam-2* (female) were used for most of the experiments. Under normal conditions, plants were grown in petri dishes containin solid 0.5× Gamborg B-5 basal medium (Phytotech #G398) at pH 5.8 with 1.2% (w/v) agar micropropagation grade (Phytotech #A296) and supplemented with 0.5% (w/v) sucrose (Merck #RDD023), under continuous LED light at 21 °C with light intensity of 150 *μ*mol/m^2^/s (Systion #SE-EGB). For spore production, plants were grown in Microbox micropropagation containers (SacO2) in long-day conditions (16 h light/8 h dark) under light supplemented with far-red light as described (Sauret-Gueto et al., 2020). For auxin treatment, media was supplemented with 3-indole acetic acid (IAA, Sigma-Aldrich #I3750) with concentrations described in the text. For MpCLE2 treatment, the media was supplemented with the corresponding synthetic peptide (KEVONGONPLHN, Activotec) with concentrations described in the text. Mp*ARF1/2* knock-in lines and transcription factor triple reporter (*pro*Mp*ERF7:mScarlet-N7, pro*Mp*BZIP7:mVenus-N7, pro*Mp*BZIP9:mTurquoise-N7)* were described before *(Das et al., 2024; Romani et al., 2024)*.

### Plant transformation

Agrobacterium tumefaciens (GV3101) cells were transformed using the freeze-thaw method and used for plant transformation of *Cam-1/2* spores as described before (Annese et al., 2025)and selected for either 20 μg/mL hygromycin (Invitrogen #10687010) or 0.5μM chlorosulfuron (Thermo Scientific #17959385). Plants were screened for positive fluorescence and at least two independent lines were selected. For expression markers, representative transgenic lines are shown, showing the consensus expression pattern out of 4-5 independent lines. In the case of bimolecular fluorescence complementation (BiFC), plants were co-transformed with two A. tumefaciens strains harbouring each plasmid and selected for hygromycin and chlorosulfuron to obtain stable transgenic lines. At least a dozen of independent transformant sporelings were analyzed for fluorescent protein reconstruction to obtain consistent results for protein-protein interactions.

Genotyping of Marchantia was done using Phire Plant Direct PCR Master Mix (Thermo Scientific #F-160S) according to manufacturer’s instructions. Mp*pin1* knock-out mutants were confirmed using previously published primers (Fisher et al., 2023) (CGAATAAATGCAGGAGACAAGGAGC & ATTGGTCCATGATGAGCGTCTTGGC).

### Macro photography

Plants were imaged using a Keyence VHX 5000 digital microscope equipped with a VH-Z20T lens (20x-200x)

### Confocal microscopy

Images were acquired using a Leica SP8X spectral confocal microscope with a 460–670 nm supercontinuum laser (70% power), two diode lasers (405 and 442 nm), four hybrid detectors, and one photomultiplier tube. Imaging used 10x air (HC PL APO 10×/0.40 CS2), 20x air (HC PL APO 20x/0.75 CS2) 40x water immersion, (HC PL APO 40x/1.10 W CORR CS2) 63x water-immersion (HC PL APO 63×/1.20 W CORR CS2) objectives. Sequential scanning (between lines) was applied for each channel, unless CFP/GFP type fluorescent protein was being imaged with chlorophyll, in which case they were collected in a single sequence. Fluorescent protein settings were most often: mTurquoise2 (152) (442 nm, 445–480 nm), mVenus (153) (515 nm, 522–560 nm), mScarlet-I (154) (569 nm, 585–625 nm), and chlorophyll autofluorescence (442 nm, 690–720 nm), but were adjusted between plant lines and tissue type to minimise collection of tissue autofluorescence signals. Images were processed FIJI.

### Optical clearing of transgenic plants

Plants were fixed with 4% (w/v) formaldehyde in PBS for 1 h under vacuum, washed with 1 mL PBS three times, then incubated with iTomei-D (#T3940, Tokyo Chemical Industry) overnight. Samples were washed again with 1 mL PBS and stained with SR2200 (157) for 6 hours, washed in 1mL PBS and destained in PBS for 1 hour. Fixed, cleared and stained tissue was placed in 100 μL 70% iohexol (#I0903, Tokyo Chemical Industry) for 30 minutes, in which they were mounted inside of two stacked Gene Frames (Thermo Fisher Scientific). Samples were covered with a glass coverslip and imaged on a Leica SP8X confocal microscope.

### Time-lapse imaging of plant tissues

Tissue fragments were placed in 2-4 stacked Gene Frames, atop ½ GB medium agar pads containing 0.5% (w/v) sucrose. 150 μL of perfluorodecalin (Sigma-Aldrich #P9900) was pipetted on top of the tissue and imaging chamber was sealed with a glass cover slip.

## Statistical analysis

Statistical analysis was performed on MS Excel and R version 4.5.1 Plots were generated using R version 4.5.1.

**SUPPLEMENTARY FIGURE 1.**
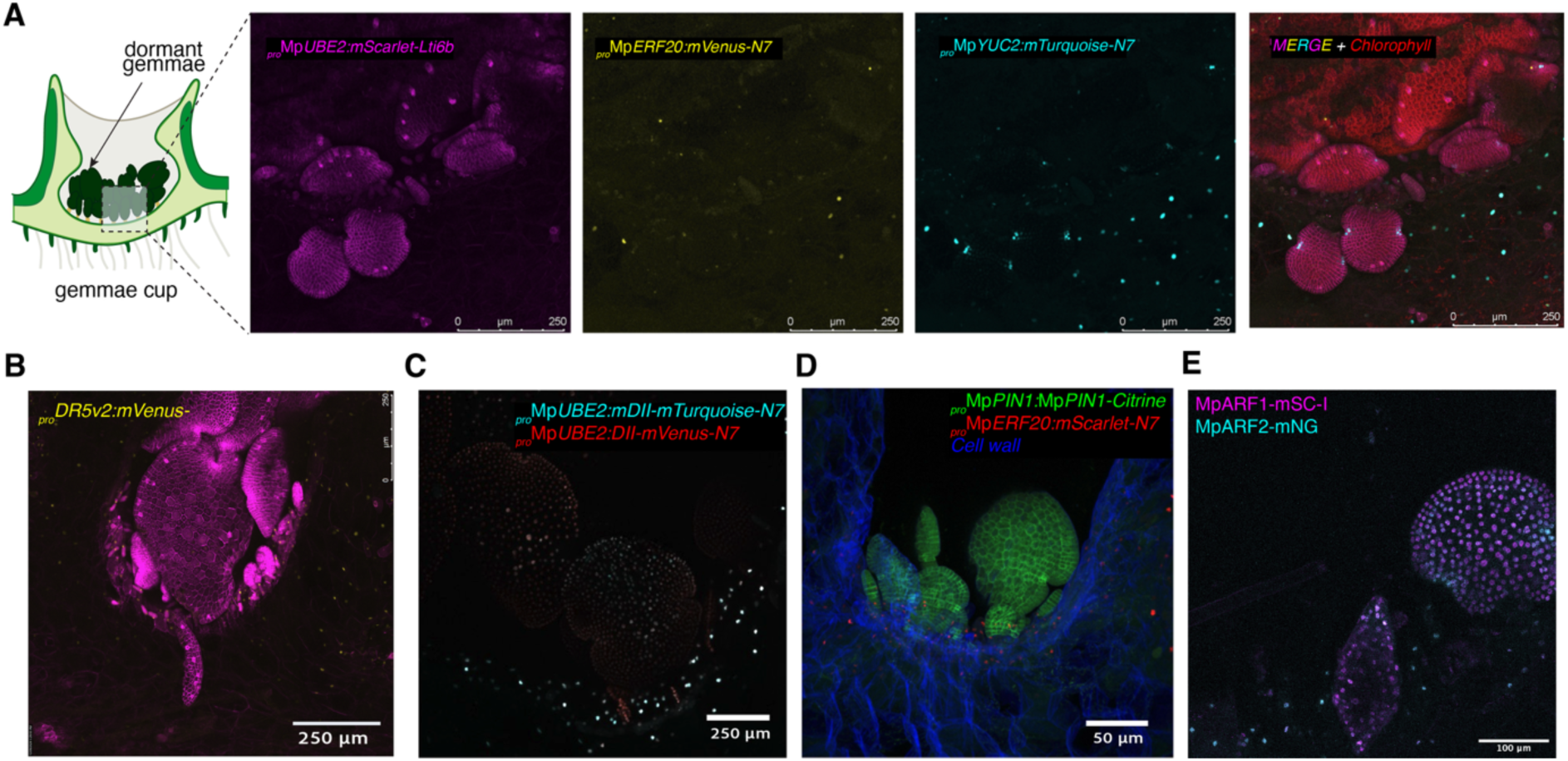
Auxin distribution and signalling in gemmae and gemma cups. (A) Confocal images of dormant gemmae within the gemma cup showing expression of a constitutive plasma membrane marker (proMpUBE2:mScarlet-Lti6b, magenta), proMpERF20:mVenus-N7 (yellow), proMpYUC2:mTurquoise-N7 (cyan), and chlorophyll autofluorescence (red). (B) Z-projection confocal image of proDR5v2:mVenus-N7 (yellow) in dormant gemmae within the cup. (C) Z-projection confocal image of the R2DII ratiometric auxin reporter (proMpUBE2:mDII-mTurquoise-N7; proMpUBE2:DII-mVenus-N7) in dormant gemmae. Red regions indicate lower auxin accumulation. (D) Confocal image of proMpPIN1:MpPIN1-Citrine (green) and proMpERF20:mScarlet-N7 (red) in gemmae within the cup, with cell wall stained with SR2200 (blue). (E) Confocal image of MpARF1-mScarlet-I (magenta) and MpARF2-mNeonGreen (cyan) knock-in lines in dormant gemmae. Scale bars = 250 µm unless otherwise indicated.

**SUPPLEMENTARY FIGURE 2.**
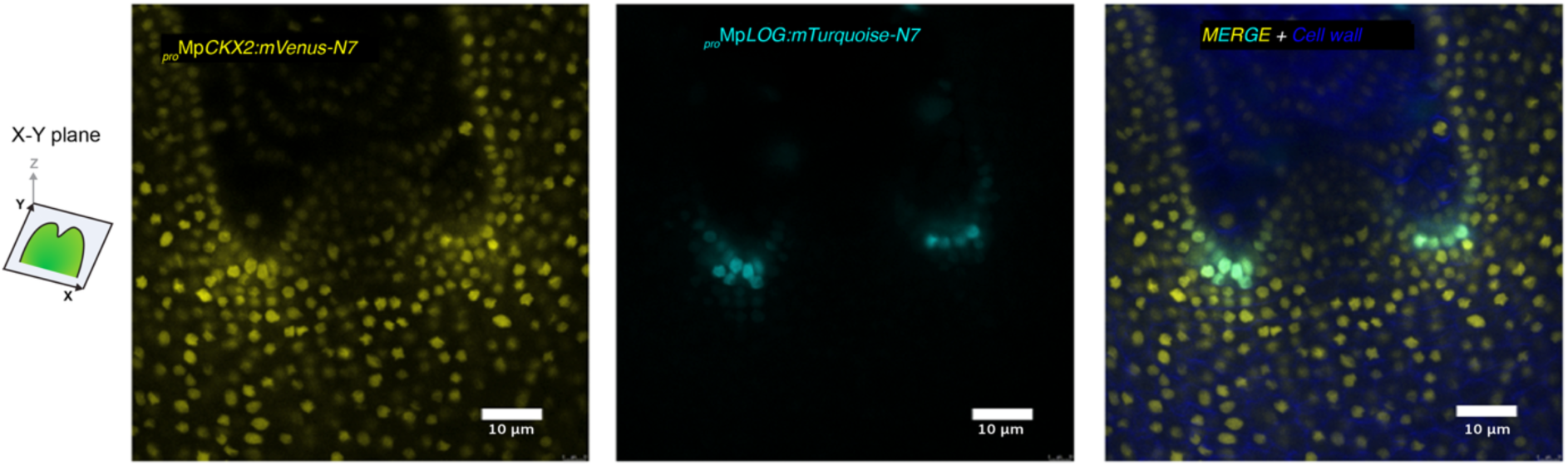
Cytokinin signalling markers in the apical meristem. X-Y plane confocal images of the apical notch of 2-week-old plants expressing proMpCKX2:mVenus-N7 (yellow, left) and proMpLOG:mTurquoise-N7 (cyan, centre), with merged image including cell wall stain (right). MpLOG expression is restricted to the apical cell and immediate sub-apical cells, consistent with an active cytokinin microenvironment in the stem cell zone. Scale bars = 10 µm.

**SUPPLEMENTARY MOVIE 1. MpPIN1 time-lapse during regeneration.** Z-stack of a time-lapse confocal image of proMpPIN1:MpPIN1–Citrine translational reporter (green) with the proMpERF20:mScarlet-N7 transcriptional reporter (red).

**SUPPLEMENTARY MOVIE 2. MpARF1/2 time-lapse during regeneration.** Z-stack of a time-lapse confocal image of MpARF1-mScarlet-I (magenta) and MpARF2-mNeonGreen (cyan) knock-in lines.

## REFERENCES

1. Alon, U. (2007). Network motifs: theory and experimental approaches. Nat Rev Genet 8, 450–461.

2. Annese, D., Romani, F., Grandellis, C., Ives, L., Frangedakis, E., Buson, F.X., Molloy, J.C., and Haseloff, J. (2025). Semi-automated workflow for high-throughput Agrobacterium-mediated plant transformation. Plant J 122, e70118.

3. Banno, H., Ikeda, Y., Niu, Q.W., and Chua, N.H. (2001). Overexpression of Arabidopsis ESR1 induces initiation of shoot regeneration. Plant Cell 13, 2609–2618.

4. Barbez, E., Kubes, M., Rolcik, J., Beziat, C., Pencik, A., Wang, B., Rosquete, M.R., Zhu, J., Dobrev, P.I., Lee, Y., Zazimalova, E., Petrasek, J., Geisler, M., Friml, J., and Kleine-Vehn, J. (2012). A novel putative auxin carrier family regulates intracellular auxin homeostasis in plants. Nature 485, 119–122.

5. Bennett, T.A., Liu, M.M., Aoyama, T., Bierfreund, N.M., Braun, M., Coudert, Y., Dennis, R.J., O’Connor, D., Wang, X.Y., White, C.D., Decker, E.L., Reski, R., and Harrison, C.J. (2014). Plasma membrane-targeted PIN proteins drive shoot development in a moss. Curr Biol 24, 2776–2785.

6. Blazquez, M.A., Nelson, D.C., and Weijers, D. (2020). Evolution of Plant Hormone Response Pathways. Annu Rev Plant Biol 71, 327–353.

7. Bowman, J.L., Kohchi, T., Yamato, K.T., Jenkins, J., Shu, S., Ishizaki, K., Yamaoka, S., Nishihama, R., Nakamura, Y., Berger, F., Adam, C., Aki, S.S., Althoff, F., Araki, T., Arteaga-Vazquez, M.A., Balasubrmanian, S., Barry, K., Bauer, D., Boehm, C.R., Briginshaw, L., Caballero-Perez, J., Catarino, B., Chen, F., Chiyoda, S., Chovatia, M., Davies, K.M., Delmans, M., Demura, T., Dierschke, T., Dolan, L., Dorantes-Acosta, A.E., Eklund, D.M., Florent, S.N., Flores-Sandoval, E., Fujiyama, A., Fukuzawa, H., Galik, B., Grimanelli, D., Grimwood, J., Grossniklaus, U., Hamada, T., Haseloff, J., Hetherington, A.J., Higo, A., Hirakawa, Y., Hundley, H.N., Ikeda, Y., Inoue, K., Inoue, S.I., Ishida, S., Jia, Q., Kakita, M., Kanazawa, T., Kawai, Y., Kawashima, T., Kennedy, M., Kinose, K., Kinoshita, T., Kohara, Y., Koide, E., Komatsu, K., Kopischke, S., Kubo, M., Kyozuka, J., Lagercrantz, U., Lin, S.S., Lindquist, E., Lipzen, A.M., Lu, C.W., De Luna, E., Martienssen, R.A., Minamino, N., Mizutani, M., Mizutani, M., Mochizuki, N., Monte, I., Mosher, R., Nagasaki, H., Nakagami, H., Naramoto, S., Nishitani, K., Ohtani, M., Okamoto, T., Okumura, M., Phillips, J., Pollak, B., Reinders, A., Rovekamp, M., Sano, R., Sawa, S., Schmid, M.W., Shirakawa, M., Solano, R., Spunde, A., Suetsugu, N., Sugano, S., Sugiyama, A., Sun, R., Suzuki, Y., Takenaka, M., Takezawa, D., Tomogane, H., Tsuzuki, M., Ueda, T., Umeda, M., Ward, J.M., Watanabe, Y., Yazaki, K., Yokoyama, R., Yoshitake, Y., Yotsui, I., Zachgo, S., and Schmutz, J. (2017). Insights into Land Plant Evolution Garnered from the Marchantia polymorpha Genome. Cell 171, 287–304 e215.

8. Cammarata, J., Roeder, A.H.K., and Scanlon, M.J. (2023). The ratio of auxin to cytokinin controls leaf development and meristem initiation in Physcomitrium patens. J Exp Bot 74, 6541–6550.

9. Das, S., de Roij, M., Bellows, S., Alvarez, M.D., Mutte, S., Kohlen, W., Farcot, E., Weijers, D., and Borst, J.W. (2024). Quantitative imaging reveals the role of MpARF proteasomal degradation during gemma germination. Plant Commun 5, 101039.

10. de Roij, M., Hernandez Garcia, J., Das, S., Borst, J.W., and Weijers, D. (2025). ARF degradation defines a deeply conserved step in auxin response. Nat Plants 11, 717–724.

11. Eklund, D.M., Ishizaki, K., Flores-Sandoval, E., Kikuchi, S., Takebayashi, Y., Tsukamoto, S., Hirakawa, Y., Nonomura, M., Kato, H., Kouno, M., Bhalerao, R.P., Lagercrantz, U., Kasahara, H., Kohchi, T., and Bowman, J.L. (2015). Auxin Produced by the Indole-3-Pyruvic Acid Pathway Regulates Development and Gemmae Dormancy in the Liverwort Marchantia polymorpha. Plant Cell 27, 1650–1669.

12. Fisher, T.J., Flores-Sandoval, E., Alvarez, J.P., and Bowman, J.L. (2023). PIN-FORMED is required for shoot phototropism/gravitropism and facilitates meristem formation in Marchantia polymorpha. New Phytol 238, 1498–1515.

13. Flores-Sandoval, E., Eklund, D.M., and Bowman, J.L. (2015). A Simple Auxin Transcriptional Response System Regulates Multiple Morphogenetic Processes in the Liverwort Marchantia polymorpha. PLoS Genet 11, e1005207.

14. Flores-Sandoval, E., Eklund, D.M., Hong, S.F., Alvarez, J.P., Fisher, T.J., Lampugnani, E.R., Golz, J.F., Vazquez-Lobo, A., Dierschke, T., Lin, S.S., and Bowman, J.L. (2018). Class C ARFs evolved before the origin of land plants and antagonize differentiation and developmental transitions in Marchantia polymorpha. New Phytol 218, 1612–1630.

15. Flores-Sandoval, E., Suzuki, H., Lazner, J.A., Briginshaw, L.N., Fisher, T.J., Romani, F., Levins, J., Hainiwa, E., Kinami, T., Imai, Y., Yumoto, E., Asahina, M., Kohchi, T., Nishihama, R., and Bowman, J.L. (2025). The B-class auxin response factor MpARF2 is essential for meristem organization in free-living plant gametophytes. Curr Biol.

16. Hata, Y., Hetherington, N., Battenberg, K., Hirota, A., Minoda, A., Hayashi, M., and Kyozuka, J. (2025). snRNA-seq analysis of the moss Physcomitriumpatens identifies a conserved cytokinin-ESR module promoting pluripotent stem cell identity. Dev Cell.

17. Heisler, M.G., Ohno, C., Das, P., Sieber, P., Reddy, G.V., Long, J.A., and Meyerowitz, E.M. (2005). Patterns of auxin transport and gene expression during primordium development revealed by live imaging of the Arabidopsis inflorescence meristem. Curr Biol 15, 1899–1911.

18. Hirakawa, Y., Fujimoto, T., Ishida, S., Uchida, N., Sawa, S., Kiyosue, T., Ishizaki, K., Nishihama, R., Kohchi, T., and Bowman, J.L. (2020). Induction of Multichotomous Branching by CLAVATA Peptide in Marchantia polymorpha. Curr Biol 30, 3833–3840 e3834.

19. Hisanaga, T., Fujimoto, S., Cui, Y., Sato, K., Sano, R., Yamaoka, S., Kohchi, T., Berger, F., and Nakajima, K. (2021). Deep evolutionary origin of gamete-directed zygote activation by KNOX/BELL transcription factors in green plants. Elife 10.

20. Ishida, S., Suzuki, H., Iwaki, A., Kawamura, S., Yamaoka, S., Kojima, M., Takebayashi, Y., Yamaguchi, K., Shigenobu, S., Sakakibara, H., Kohchi, T., and Nishihama, R. (2022). Diminished Auxin Signaling Triggers Cellular Reprogramming by Inducing a Regeneration Factor in the Liverwort Marchantia polymorpha. Plant Cell Physiol 63, 384–400.

21. Kaplan, S., Bren, A., Dekel, E., and Alon, U. (2008). The incoherent feed-forward loop can generate non-monotonic input functions for genes. Mol Syst Biol 4, 203.

22. Kato, H., Ishizaki, K., Kouno, M., Shirakawa, M., Bowman, J.L., Nishihama, R., and Kohchi, T. (2015). Auxin-Mediated Transcriptional System with a Minimal Set of Components Is Critical for Morphogenesis through the Life Cycle in Marchantia polymorpha. PLoS Genet 11, e1005084.

23. Kato, H., Kouno, M., Takeda, M., Suzuki, H., Ishizaki, K., Nishihama, R., and Kohchi, T. (2017). The Roles of the Sole Activator-Type Auxin Response Factor in Pattern Formation of Marchantia polymorpha. Plant Cell Physiol 58, 1642–1651.

24. Kato, H., Mutte, S.K., Suzuki, H., Crespo, I., Das, S., Radoeva, T., Fontana, M., Yoshitake, Y., Hainiwa, E., van den Berg, W., Lindhoud, S., Ishizaki, K., Hohlbein, J., Borst, J.W., Boer, D.R., Nishihama, R., Kohchi, T., and Weijers, D. (2020). Design principles of a minimal auxin response system. Nat Plants 6, 473–482.

25. Kawamura, S., Romani, F., Yagura, M., Mochizuki, T., Sakamoto, M., Yamaoka, S., Nishihama, R., Nakamura, Y., Yamato, K.T., Bowman, J.L., Kohchi, T., and Tanizawa, Y. (2022). MarpolBase Expression: A Web-Based, Comprehensive Platform for Visualization and Analysis of Transcriptomes in the Liverwort Marchantia polymorpha. Plant Cell Physiol 63, 1745–1755.

26. Kim, D., Paggi, J.M., Park, C., Bennett, C., and Salzberg, S.L. (2019). Graph-based genome alignment and genotyping with HISAT2 and HISAT-genotype. Nat Biotechnol 37, 907–915.

27. Kirch, T., Simon, R., Grunewald, M., and Werr, W. (2003). The DORNROSCHEN/ENHANCER OF SHOOT REGENERATION1 gene of Arabidopsis acts in the control of meristem ccll fate and lateral organ development. Plant Cell 15, 694–705.

28. Komatsu, A., Fujibayashi, M., Kumagai, K., Suzuki, H., Hata, Y., Takebayashi, Y., Kojima, M., Sakakibara, H., and Kyozuka, J. (2025). KAI2-dependent signaling controls vegetative reproduction in Marchantia polymorpha through activation of LOG-mediated cytokinin synthesis (14). Nat Commun 16, 1263.

29. Liao, C.Y., Smet, W., Brunoud, G., Yoshida, S., Vernoux, T., and Weijers, D. (2015). Reporters for sensitive and quantitative measurement of auxin response. Nat Methods 12, 207–210, 202 p following 210.

30. Ma, Y., Miotk, A., Sutikovic, Z., Ermakova, O., Wenzl, C., Medzihradszky, A., Gaillochet, C., Forner, J., Utan, G., Brackmann, K., Galvan-Ampudia, C.S., Vernoux, T., Greb, T., and Lohmann, J.U. (2019). WUSCHEL acts as an auxin response rheostat to maintain apical stem cells in Arabidopsis. Nat Commun 10, 5093.

31. Mangan, S., and Alon, U. (2003). Structure and function of the feed-forward loop network motif. Proc Natl Acad Sci U S A 100, 11980–11985.

32. Mellor, N., Band, L.R., Pencik, A., Novak, O., Rashed, A., Holman, T., Wilson, M.H., Voss, U., Bishopp, A., King, J.R., Ljung, K., Bennett, M.J., and Owen, M.R. (2016). Dynamic regulation of auxin oxidase and conjugating enzymes AtDAO1 and GH3 modulates auxin homeostasis. Proc Natl Acad Sci U S A 113, 11022–11027.

33. Moreno-Risueno, M.A., Van Norman, J.M., Moreno, A., Zhang, J., Ahnert, S.E., and Benfey, P.N. (2010). Oscillating gene expression determines competence for periodic Arabidopsis root branching. Science 329, 1306–1311.

34. Nishihama, R., Ishizaki, K., Hosaka, M., Matsuda, Y., Kubota, A., and Kohchi, T. (2015). Phytochrome-mediated regulation of cell division and growth during regeneration and sporeling development in the liverwort Marchantia polymorpha. J Plant Res 128, 407–421.

35. Patron, N.J., Orzaez, D., Marillonnet, S., Warzecha, H., Matthewman, C., Youles, M., Raitskin, O., Leveau, A., Farre, G., Rogers, C., Smith, A., Hibberd, J., Webb, A.A., Locke, J., Schornack, S., Ajioka, J., Baulcombe, D.C., Zipfel, C., Kamoun, S., Jones, J.D., Kuhn, H., Robatzek, S., Van Esse, H.P., Sanders, D., Oldroyd, G., Martin, C., Field, R., O’Connor, S., Fox, S., Wulff, B., Miller, B., Breakspear, A., Radhakrishnan, G., Delaux, P.M., Loque, D., Granell, A., Tissier, A., Shih, P., Brutnell, T.P., Quick, W.P., Rischer, H., Fraser, P.D., Aharoni, A., Raines, C., South, P.F., Ane, J.M., Hamberger, B.R., Langdale, J., Stougaard, J., Bouwmeester, H., Udvardi, M., Murray, J.A., Ntoukakis, V., Schafer, P., Denby, K., Edwards, K.J., Osbourn, A., and Haseloff, J. (2015). Standards for plant synthetic biology: a common syntax for exchange of DNA parts. New Phytol 208, 13–19.

36. Peret, B., Swarup, K., Ferguson, A., Seth, M., Yang, Y., Dhondt, S., James, N., Casimiro, I., Perry, P., Syed, A., Yang, H., Reemmer, J., Venison, E., Howells, C., Perez-Amador, M.A., Yun, J., Alonso, J., Beemster, G.T., Laplaze, L., Murphy, A., Bennett, M.J., Nielsen, E., and Swarup, R. (2012). AUX/LAX genes encode a family of auxin influx transporters that perform distinct functions during Arabidopsis development. Plant Cell 24, 2874–2885.

37. Ranocha, P., Dima, O., Nagy, R., Felten, J., Corratge-Faillie, C., Novak, O., Morreel, K., Lacombe, B., Martinez, Y., Pfrunder, S., Jin, X., Renou, J.P., Thibaud, J.B., Ljung, K., Fischer, U., Martinoia, E., Boerjan, W., and Goffner, D. (2013). Arabidopsis WAT1 is a vacuolar auxin transport facilitator required for auxin homoeostasis. Nat Commun 4, 2625.

38. Romani, F., and Moreno, J.E. (2021). Molecular mechanisms involved in functional macroevolution of plant transcription factors. New Phytol 230, 1345–1353.

39. Romani, F., Bonter, I., Rebmann, M., Takahashi, G., Guzman-Chavez, F., De Batte, F., Hirakawa, Y., and Haseloff, J. (2026). A simple cell-cycle control system in Marchantia polymorpha provides a framework for understanding plant cell proliferation. Plant Cell.

40. Romani, F., Sauret-Gueto, S., Rebmann, M., Annese, D., Bonter, I., Tomaselli, M., Dierschke, T., Delmans, M., Frangedakis, E., Silvestri, L., Rever, J., Bowman, J.L., Romani, I., and Haseloff, J. (2024). The landscape of transcription factor promoter activity during vegetative development in Marchantia. Plant Cell 36, 2140–2159.

41. Sakamoto, Y., Ishimoto, A., Sakai, Y., Sato, M., Nishihama, R., Abe, K., Sano, Y., Furuichi, T., Tsuji, H., Kohchi, T., and Matsunaga, S. (2022). Improved clearing method contributes to deep imaging of plant organs. Commun Biol 5, 12.

42. Sauret-Gueto, S., Frangedakis, E., Silvestri, L., Rebmann, M., Tomaselli, M., Markel, K., Delmans, M., West, A., Patron, N.J., and Haseloff, J. (2020). Systematic Tools for Reprogramming Plant Gene Expression in a Simple Model, Marchantia polymorpha. ACS Synth Biol 9, 864–882.

43. Spencer, V., Wallner, E.S., Jandrasits, K., Edelbacher, N., Mosiolek, M., and Dolan, L. (2024). Three-dimensional anatomy and dorsoventral asymmetry of the mature Marchantia polymorpha meristem develops from a symmetrical gemma meristem. Development 151.

44. Suzuki, H., Harrison, C.J., Shimamura, M., Kohchi, T., and Nishihama, R. (2020). Positional cues regulate dorsal organ formation in the liverwort Marchantia polymorpha. J Plant Res 133, 311–321.

45. Suzuki, H., Kato, H., Iwano, M., Nishihama, R., and Kohchi, T. (2022). Auxin signaling is essential for organogenesis but not for cell survival in the liverwort Marchantia polymorpha. Plant Cell.

46. Tang, H., Smoljan, A., Zou, M., Zhang, Y., Lu, K.J., and Friml, J. (2026). The miniW Domain Directs Polarized Membrane Localization of Non-Canonical PINs in Marchantia polymorpha. Plant Cell Environ 49, 1505–1508.

47. Thelander, M., Landberg, K., and Sundberg, E. (2019). Minimal auxin sensing levels in vegetative moss stem cells revealed by a ratiometric reporter. New Phytol 224, 775–788.

48. Tse, S.W., Annese, D., Romani, F., Guzman-Chavez, F., Bonter, I., Forestier, E., Frangedakis, E., and Haseloff, J. (2024). Optimizing Promoters and Subcellular Localization for Constitutive Transgene Expression in Marchantia polymorpha. Plant Cell Physiol 65, 1298–1309.

49. Ulmasov, T., Murfett, J., Hagen, G., and Guilfoyle, T.J. (1997). Aux/IAA proteins repress expression of reporter genes containing natural and highly active synthetic auxin response elements. Plant Cell 9, 1963–1971.

50. Vernoux, T., Besnard, F., and Traas, J. (2010). Auxin at the shoot apical meristem. Cold Spring Harb Perspect Biol 2, a001487.

51. Viaene, T., Landberg, K., Thelander, M., Medvecka, E., Pederson, E., Feraru, E., Cooper, E.D., Karimi, M., Delwiche, C.F., Ljung, K., Geisler, M., Sundberg, E., and Friml, J. (2014). Directional auxin transport mechanisms in early diverging land plants. Curr Biol 24, 2786–2791.

52. Wallner, E.S., and Dolan, L. (2024). Reproducibly oriented cell divisions pattern the prothallus to set up dorsoventrality and de novo meristem formation in Marchantia polymorpha. Curr Biol 34, 4357–4367 e4354.

53. Wallner, E.S., Edelbacher, N., and Dolan, L. (2026). De novo meristem development in Marchantia polymorpha requires light and an apical auxin signaling minimum. Curr Biol 36, 278–289 e275.

54. Zhang, C., Tsoi, R., Wu, F., and You, L. (2016). Processing Oscillatory Signals by Incoherent Feedforward Loops. PLoS Comput Biol 12, e1005101.

